# Transcriptome analysis reveals regulatory networks and hub genes in flavonoid metabolism of *Rosa roxburghii*

**DOI:** 10.1101/2021.04.06.438570

**Authors:** Xiaolong Huang, Zongmin Wu, Yanjing Liu, Yin Yi, Huiqing Yan, Qiaohong Li

## Abstract

*Rosa roxburghii* Tratt, the most popular fruit that blooms in the southwest of China, is rich in flavonoids. However, the regulatory network and critical genes involved in the metabolism of flavonoid compounds in *R. roxburghii* are still unknown. In this study, we revealed that flavonoid, anthocyanin and catechin accumulated at different levels in various tissues of *R. roxburghii*. We further obtained and analyzed differentially expressed genes (DEGs) involved in flavonoid metabolism from five samples of *R. roxburghii* by transcriptome sequencing. A total of 1 130 DEGs were identified, including 166 flavonoid pathway biosynthesis genes, 622 transcription factors, 301 transporters, and 221 cytochrome P450 proteins. A weighted gene co-expression network analysis (WGCNA) of the DEGs uncovered different co-expression networks. In terms of biosynthesis enzymes, cytochrome P450 CYP749A22 and CYP72A219 were highlighted in regulation flavonoids content. Anthocyanin 3-O-glucosyltransferase and F3’H were the top two critical enzymes for anthocyanin content. In contrast, caffeic acid 3-O-methyltransferase, 4-coumarate-CoA ligase, and shikimate O-hydroxycinnamoyl transferase were essential for catechin accumulation. Additionally, the eigengene network of the “black” module had high correlations with total flavonoid (r= 0.9, p=5e-06). There were 26 eigengenes in the “black” module, including six flavonoid biosynthesis, 14 TFs and six transporters. Among them, besides cytochrome P450 proteins (*DN136557_c0_g1*, *DN135573_c0_g1* and *DN145971_c4_g1*), isoflavone-hydroxylase (*DN143321_c3_g1*) was crucial for total flavonoids content based on the high degree of connectivity. The transcription factors *RrWRKY45* (*DN142829_c1_g5*), *RrTCP20* (*DN146443_c1_g1*) and *RrERF118* (*DN141507_c3_g2*) were significantly correlated with flavonoids in *R. roxburghii*. The present transcriptomic and biochemical data on metabolites should encourage further investigation on functional genomics and breeding of *R. roxburghii* with strong pharmaceutical potential.

## Introduction

*Rosa roxburghii* Tratt, a deciduous shrub and Chinese ethnomedicine which belongs to the *Rosacea* family, is commercially cultivated, and consumed in China. Flavonoids extracted from *R. roxburghii* have high antioxidant potential and beneficial health properties. They are reported to exert antioxidants functions, scavenge free radicles, reduce high cholesterol, and have antimutagenic properties [1]. Besides, they offer protection against radiation-induced apoptosis and inflammation [2–4]. Previous studies show that *R. roxburghii* is rich in flavonoids, especially highly accumulates in fruits with a purity of 73.85% [4–6]. Nevertheless, little information is available regarding the exact molecular mechanism underlying flavonoids accumulation in *R. roxburghii*.

Flavonoids, a kind of polyphenolic metabolites, play multiple physiological roles in plant development and are known for wide distribution in plants, even in different organs [7]. Flavonoids comprise many compounds, including anthocyanins, isoflavones, flavanones, flavanols, catechin, and a series of phenolic acid derivatives synthesized through the phenylpropanoid and different branched sub pathways [8, 9]. The precursors of most flavonoids are malonyl-CoA and p-coumaroyl-CoA and that the biosynthesis begins in the phenylpropanoid pathway. Some key biosynthesis enzymes directly regulate flavonoid accumulation. The deletion of chalcone synthase (CHS) induced no flavonoid accumulation in *Arabidopsis tt4* (*transparent testa4*) plants [10]. In addition to flavonoid pathway-related enzymes, cytochrome P450 proteins, a large family of heme-containing monooxygenases, catalyze diverse types of chemical reactions and play an essential role in flavonoid metabolism in general [11]. The generations of p-Cinnamoyl-CoA and cinnamic acid 4-hydroxylase are catalyzed by CYP73A [12]. CYP98 is responsible for an enzyme hydroxylating the 3’-position of the phenolic ring [13]. Kaempferol can be transformed into quercetin through catalysis by CYP75A [14]. Cytochrome P450 proteins are also involved in the biosynthesis of flavones [CYP93B, (2S)-flavanone 2-hydroxylase and flavone synthase II], and leguminous isoflavonoid phytoalexins [YP93A, 3,9-dihydroxypterocarpan 6a-hydroxylase; CYP93C, 2-hydroxyisoflavanone synthase (IFS)] [15].

The types and contents of flavonoids depend not only on the flavonoid biosynthesis enzymes but also depends on the bio-modification, transportation processes and transcription factors [16]. H^+^- ATPases are the primary driving force to transport flavonoids [17]. ABC (ATP-binding cassette) and MATE (multidrug and toxic compound extrusion protein) transporters are also assumed to play essential roles in the sequestration of flavonoids into the vacuole [18–20]. Moreover, MVT (membrane vesicle-mediated) transport, requiring VSR (vacuolar sorting receptor) and SNARE (soluble N-ethylmaleimide sensitive factor attachment protein receptors) proteins, involves flavonoid accumulation [21]. GST (Glutathione S-transferase) catalyzes the conjugation of the tripeptide glutathione as flavonoid binding proteins and is responsible for transport [22]. Besides, numerous transcription factors (TFs) influence the biosynthesis of flavonoid glycosides in plants [23]. MYBs (v-myb avian myeloblastosis viral oncogene homolog) activate the structural genes from the flavonoid pathway branch leading to the generation of flavonoid [24]. MYBs interact with bHLH (basic helix-loop-helix) and WD40 proteins together to form an MYB-bHLH-WD40 complex in regulating flavonoid metabolism [25–27]. Additionally, NAC (NAM, ATAF1/2, and CUC2) was reported to up-regulate the transcript levels expression of genes associated with the biosynthesis of flavonoids. The expression of some flavonoid biosynthesis genes and the content of anthocyanins were significantly increased in *AtNAC* overexpression transgenic plants and reduced when *AtNAC* was knockout, implying that *AtNAC* influences the flavonoid accumulation [28].

Given so many factors could influence flavonoid compounds, it is essential to reveal the critical genes which play important roles in determining the total flavonoids content in *R. roxburghii*. An *R* analytical package named weighted gene co-expression network analysis (WGCNA) is a systems biology approach aimed at understanding networks between genes instead of any individual genes [29, 30]. The highest-degree nodes are often called ‘hubs’, thought to serve specific purposes in their networks. Therefore, WGCNA has been widely used to determine different gene modules of highly correlated genes and provide some highly connected hub genes [31, 32]. The previous paper showed ten transporters and seven TFs were critical genes in anthocyanin accumulation of *Lycium ruthenicum* Murray [33]. Twenty-four genes encoding proteins putatively associated with anthocyanin regulation, biosynthesis, and transport were critical for anthocyanin in an apple (*Malus* × *domestica*) yellow fruit somatic mutation [34].

Considered the distributions of flavonoid vary differently among various tissues, the transcriptome of the *R. roxburghii* stem, leaf, flower, young fruit, and mature fruit has been conducted using the Illumina HiSeq 3000 platform [35]. In this study, to obtain insights into the changes of flavonoid metabolites and transcriptional regulation in *R. roxburghii*, differentially expressed flavonoid genes were identified. Furthermore, we obtained flavonoid related co-expression gene modules and screened some critical genes involved in flavonoid synthesis by WGCNA. This study unravels the key genes that influenced flavonoids content and further refines the *R. roxburghii* flavonoid regulatory network.

## Materials and methods

### Plant Materials and sampling

Seedlings of *R. roxburghii* were planted under natural conditions in the Guizhou Normal University, Guizhou province, China (Guiyang N 26°42.408’; E 106°67.353’). Tissues were collected, including stems, leaves, flowers, fruits at young (50 DAF, days after flowering) and mature development (120 DAF). The five samples were snap-frozen in liquid nitrogen, then mechanically ground into a fine powder and finally kept at −80°C for subsequent experiments.

### Extraction and Quantification of total flavonoid, anthocyanin, and catechin

Samples (10 mg) were extracted using 500 μL 80 % methanol, processing with ultrasonic treatment for one h followed by incubation at 4 °C for 18 h. The homogenates were centrifuged at 12 000 rpm for 15 min and the supernatant was stored for measuring [36]. A total of 50 μL the extraction was transferred into 1 mL tube with 450 μL ddH_2_O, 30 μL NaNO_2_ was added before shaking, and the reaction mixture was left for 5min.Then, 30μL 10% Al(NO_3_)_3_ was added mixed for10 min at room temperature. Then, 200 μL of 1 mol/L NaOH was added to the tube, followed by the addition of ddH_2_O up to a volume of 1 mL. The absorbance of the mixtures was measured at 500 nm, and contents of the total flavonoids of five samples were calculated with commercial rutin (No#380709, Sigma-Aldrich) as standard. The total flavonoids were expressed as mg/100 g of fresh weight (FW) [5].

The sample solution was filtered by 0.45 μm organic membranes. An amount of 20 μL was injected to determine the content of the anthocyanin and catechin. The HPLC analysis of anthocyanin was performed by adding 3% (v/v) formic acid to the methanolic extract. Dried Delphinidin 3,5-diglucoside chloride (No#PHL89626, Supelco) and catechin (No#43412, Supelco) standard materials were used to make regression equations to measure the content of catechin in samples. Chromatographic conditions-A C18 column (4.6 mm × 200 mm, 5 μm, Waters Corporation) was used. The mobile phase consisted of A: 5% (v/v) formic acid in water; solvent B: 5% (v/v) formic acid in acetonitrile. The anthocyanin was separated by starting with 100%A with a linear gradient to 25% B over 20 min, ramping to 80% B over 2 and 3 min to re-equilibrate to initial in acetonitrile. The absorption was evaluated at 520 nm [37]. The catechin was separated by the solution consisted of methanol (solvent A) and pure water (solvent B), and the flow rate was 1.0 mL/min. The optimal gradient program started with 15 % methanol and was kept in the 0-20 min period, while 25 % methanol was kept in the 20-33 min period. The column oven temperature was maintained at 30 °C. The eluent was passed through a photodiode array (PDA) detector (Waters Corporation) and the detection wavelength was evaluated at 270 nm according to the method reported by Hao et al. [5]

### RNA-Seq data, gene level quantification, analysis of differentially expressed genes (DEGs) and annotation

Five different samples were applied to high-throughput Illumina sequencing and been published [35]. Tissue collection and preparation were performed as previously described [35]. The reads were *De novo* assembly using the Trinity with default settings based on the de Bruijn graph algorithm [31]. All clean reads generated by Illumina sequencing have been deposited, being publicly available and can be readily queried in NCBI Gene Expression Omnibus (GEO) database under the accession number GSE122014. FPKM (fragments per kilo base of transcript per million mapped reads) value of each gene was calculated using cufflinks [38], and the read counts were determined by htseq-count [39]. The data were normalized the gene expression using the DESeq2 protocol [40]. A corrected adjusted P-value of FDR≤ 0.01 and abs [log2 (Fold change)] ≥ 1 were performed as the threshold to identify the significance of DEGs based on their FPKM values. Genes with FPKM lower than 0.5 were considered not expressed [29]. To identify the DEGs related to the flavonoid, we performed BLASTx alignment with an E-value ≤ 10^-5^ with different databases, including the Nr (non-redundant) protein database (https://www.ncbi.nlm.nih.gov/), and KEGG (Kyoto Encyclopedia of Genes and Genomes, http://www.genome.jp/kegg/) [41]. The GO (Gene Ontology) was determined by Blast2GO and WEGO software were used to analyze the GO functional classification [42, 43].

### Co-expression network analysis with WGCNA

Functional unigenes were annotated by blastx against. The highly co-expressed gene modules were based on FPKM values of DEGs associated with flavonoid biosynthesis, transport, transcription factors and cytochrome P450 using an R package named weighted gene co-expression network analysis (WGCNA) [30]. The WGCNA network and module were conducted using an unsigned type of topological overlap matrix (TOM). The calculation parameters “soft thresholding power” = 2 and “merge Cut Height” = 0.5 were selected to analysis of the DEGs. Finally, the module eigengene value was calculated to evaluate the relationships among modules, total flavonoid, anthocyanin and catechin content in the five samples. of modules with each tissue type.

### Identification of hub genes

The kME was determined as the Pearson correlation coefficient between each gene. The module eigengene was used to evaluate the association of the module with total flavonoids contents. The most significant module (‘black’) of genes with WGCNA edge weight >0.80 was represented using Cytoscape 7.1. The hub genes of this module were screened based on the number of edges (degree of connectivity) associated with the corresponding nodes within the network. The selection parameters were set as the top nodes ranked with the maximum number of edges in the module.

### RNA isolation and quantitative real-time PCR (qRT-PCR) analysis

Tissues used for RT-qPCR were the same batch of plants as used for RNA-seq. Firstly, RNA was extracted from five samples using the Trizol method (Takara, Japan) with the addition of RNAiso-mate to remove polysaccharides and polyphenol substances. Then the purified RNA was reversed transcribed using RT-PCR Kit (TaKaRa, Japan) with an oligo dT-adaptor primer in accordance with the manufacturer’s instructions. An SYBR Premix Ex Taq kit (Takara) was used to quantitative real-time RT-PCR. Amplification was carried out according to the following cycling parameters: denaturing for 10 min at 95 °C, 40 cycles of denaturation at 95 °C for 15 s, annealing for 30 s at 55 °C, and extension at 72 °C for 30 s. The primer pairs were designed using Primer Premier 6 and listed in Table S7. To ensure the reproducibility of results, we carried out quantitative PCR analysis in triplicate for each sample. β-actin was used as an internal control. The value of a 2^—ΔΔCt^ method was used to assess the relative gene expression.

### Statistical analysis

The data were expressed as mean ± standard deviation (SD) and analyzed using analysis of variance (ANOVA) and SPSS (Statistical Package for Social Sciences) Statistical 20.0. Statistical significance was set at p < 0.05.

## Results

### Quantification of Total Flavonoids, Anthocyanin and catechin in different tissues

The previous studies proved that the leaves and fruits of *R. roxburghii* are rich in total flavonoids and catechin [4, 5, 44]. In this study, the tissues from stems, leaves, flowers, young fruits, and mature fruits were further used to investigate the distribution of total flavonoids in *R. roxburghii*. The flavonoid was present in all tissues (Fig. 1). Their content was highest in mature fruits (237.03 mg/100 g FW), followed by young fruits, flowers, and leaves, with the contents between 70.27-175.87 mg/100 g FW. Less than 45 mg/100 g FW in stems of mature seedlings. However, anthocyanins were highly accumulated in flowers of *R. roxburghii*. The content of anthocyanin in flowers was 5.63 mg/100 g FW, followed by the other four tissues, ranging from 2.27 - 0.90 mg/100 g FW. Moreover, we determined the content of catechin in five samples. The values varied from 8.47 to 1.10 mg/100 g FW with a descending order of leaves, mature fruits, young fruits, flowers, and stems, confirming that the leaves were highly rich in catechin. These results indicated that total flavonoids and specific components of flavonoids could accumulate in various tissues of *R. roxburghii* at relatively high concentrations. Therefore, to decipher the hub genes which regulated these flavonoids, especially the accumulation of total flavonoid, we performed transcriptome sequencing from various tissues in this study.

**Fig. 1.**
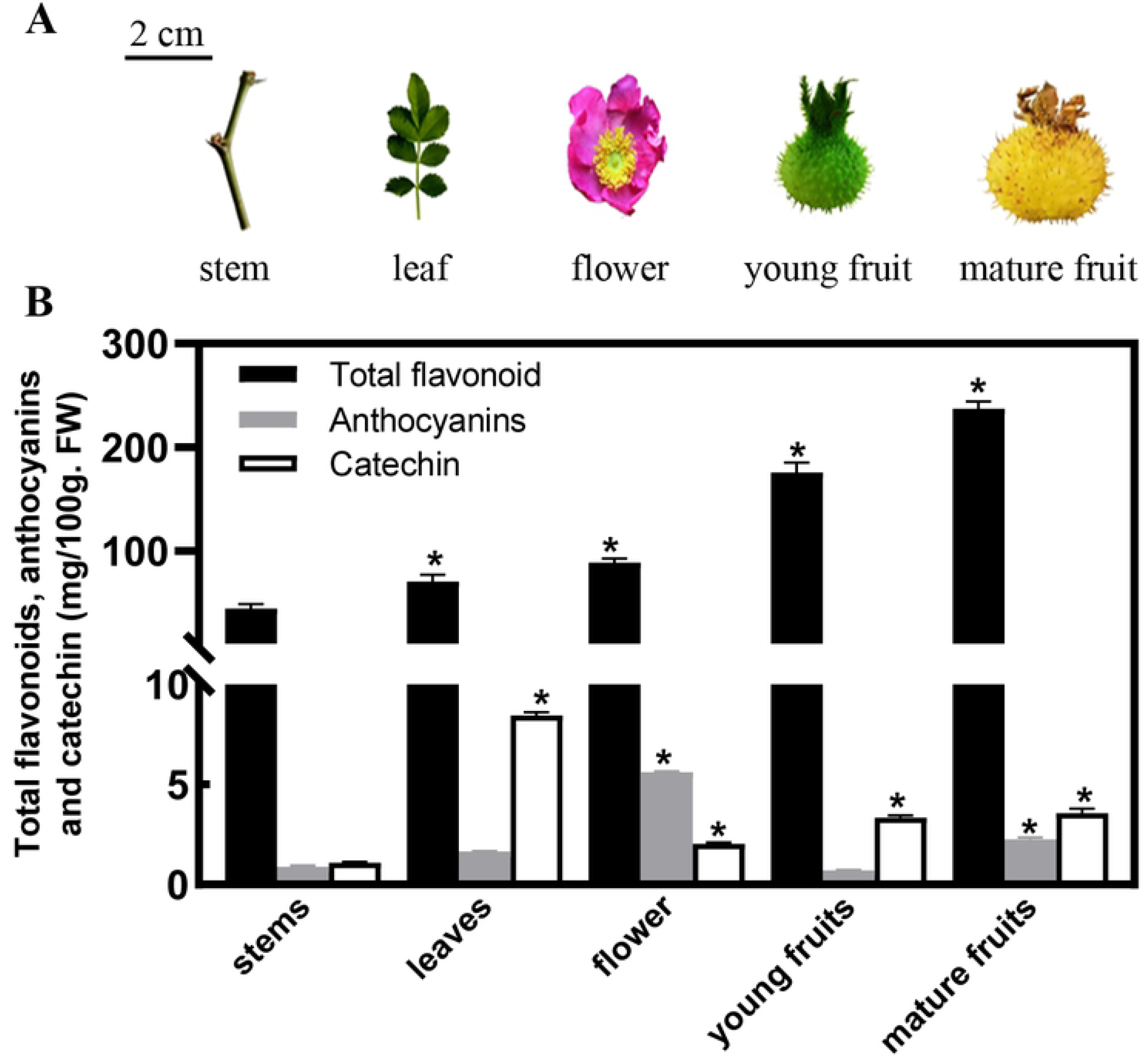
Contents of total flavonoid, anthocyanin, catechin in different tissues of *R. roxburghii*. The horizontal axis indicates different tissues, including stem, leaf, flower, young fruit, and mature fruit. The contents were expressed as mg/100 g of the fresh weight. Vertical bars represented the means ± standard deviation of three separate experiments.

### Analysis of differentially expressed genes

To preliminarily explore gene expression and then analyze the genes that regulated flavonoids in different tissues, we determined a total of 25 449 DEGs between any two tissues based on a corrected adjusted P-value of FDR≤ 0.01 and abs [log2 (Fold change)] ≥ 1. We searched for the genes related to flavonoids content and regulated genes among the DEGs based on the GO enrichment and KEGG pathway analysis of the unigenes (Fig. S1). The GO showed a total of 52.97% DEGs were involved in the metabolic process (Fig. S1A). Moreover, the KEGG enrichment displayed that phenylpropanoid synthesis (44), flavonoid biosynthesis (10), anthocyanin biosynthesis (3), flavone and flavanol biosynthesis (3) were classified, suggesting that the secondary metabolic processes are active pathways in *R. roxburghii* development (Fig. S1B).

### DEGs involved in flavonoid biosynthesis pathway

Based on GO, KEGG and the Nr library comparison using BLASTx alignment, we evaluated enzyme-encoding genes of the flavonoid pathways (Table S1). The analysis of transcriptome data uncovered that 166 key candidates exerted direct influence over twenty enzymes that were known to be involved in flavonoids KEGG biosynthesis pathways. We identified multiple transcripts encoding almost all known enzymes involved in flavonoid biosynthesis through the annotated *R. roxburghii* transcriptome. A brief outline has been shown in Fig. 2. Firstly, *RrPAL* (phenylalanine ammonia-lyase, 2 DEGs) catalyzes phenylalanine into cinnamic acid. Subsequently, *Rr4CL* (4-coumarate CoA ligase, 15 DEGs), *RrCHS* (chalcone synthase, 6 DEGs) and *RrCHI* (chalcone isomerase, 1 DEG) catalyze the cinnamic acid into naringenin or other flavonoids; flavone and flavanol biosynthesis can be produced by *RrANS* (anthocyanidin synthase, 1 DEGs). All expression levels of DEGs showed differences in five samples. *RrFLS* (flavanol synthase, 12 DEGs) and *RrDFR* (dihydroflavanol 4-reductase, 1 DEGs) take a large proportion of total flavonoids in plants [45]. Their expressions were significantly up-regulated in mature fruits showed in Fig. 2.

**Fig. 2.**
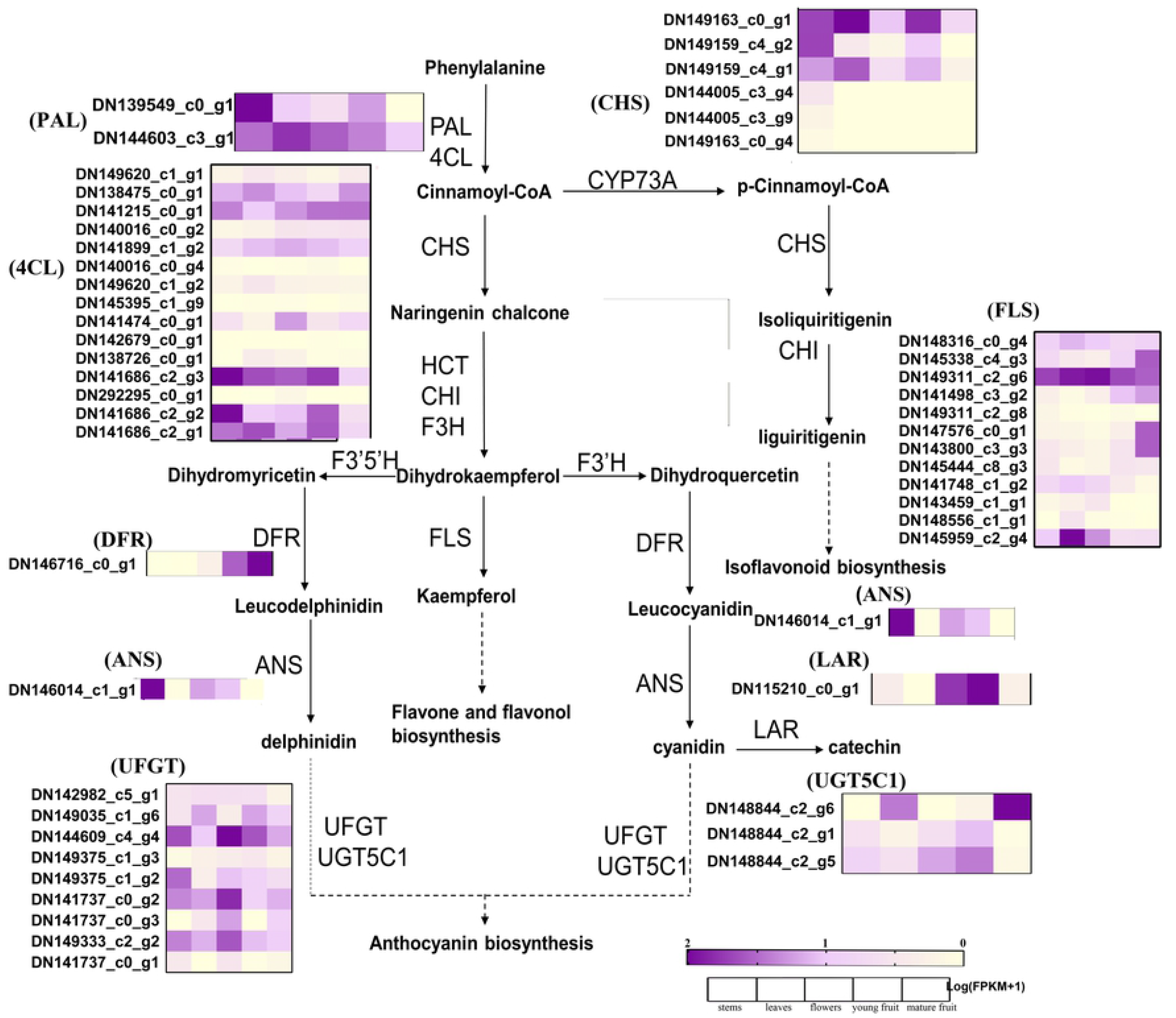
Simplified scheme and heatmap of flavonoid biosynthesis pathway genes in *R. roxburghii* based on KEGG pathway. Abbreviation: PAL phenylalanine ammonia-lyase; 4CL, 4-coumarate-CoA ligase; CHS, chalcone synthase; CHI, chalcone isomerase; ANS, anthocyanidin synthase; FLS, flavanol synthase; DFR, dihydroflavonol-4-reductase; LAR, leucoanthocyanidin reductase; ANR, anthocyanidin reductase. UFGT, UDP-glucose flavonoid 3-O-glucosyltransferase; UGT75C1, anthocyanidin 3-O-glucoside 5-O-glucosyltransferase. Arrows represent enzyme reactions; enzymes are shown next to the arrows. The color scale represents the log transformed FPKM+1 value. Purple indicates high expression, and yellow indicates low expression.

Some structures of flavonoids are very unstable. The metabolites are glycosylated, methylated and acylated before transport to vacuole storage. Thus, glycosyltransferases (*RrUFGT*, and *RrUGT75C1*) were identified. *RrUFGTs* were highly expressed in flowers (*DN144609_c4_g4*, *DN141737_c0_g2* and *DN149333_c2_g2*), while, *RrUGT75C1s-related* genes (*DN148844_c2_g6*) were highly expressed in mature fruits.

### DEGs involved in flavonoid transport and accumulation

The contents of flavonoids depend not only on the flavonoid biosynthesis pathway but also on the transportation processes [14]. According to the reported distinctive transporters [46], we determined 301 DEGs encoding candidate transporters (Table S2). The expression levels of ABC transporters were claimed to sequestration of flavonoids into the vacuole. The relations between the expression of ABC transporter genes and flavonoids content were studied [47]. In this study, 192 DEGs encoding ABC transporters were identified. The role of GST in plants was to be the carrier of flavonoids to transport [48]. A total of 80 DEGs encoding GST. Besides, 14 DEGs encoding MATE, 13 DEGs encoding SNARE, and two encoding H^+^-ATPases were identified (Table S3). The expression levels of transporters, including ABC transporter, SNARE, GST, and MATE were shown in Figure 3A. However, most of the candidate ABC transporters and GST displayed no expression in five samples.

**Fig.3.**
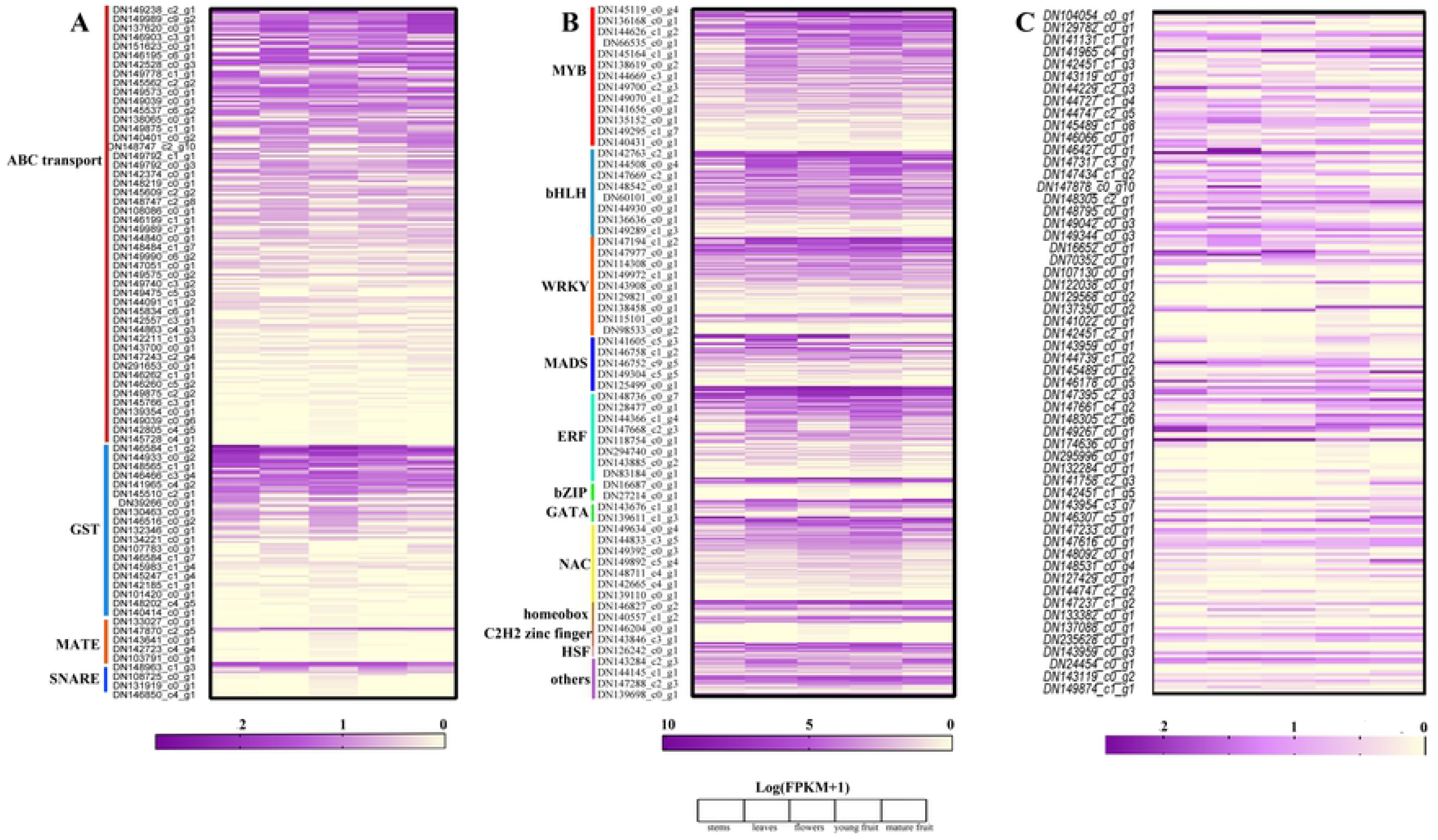
The expression levels of genes annotated with (A) transporters, (B) transcription factors and (C) P450 in five samples.

### DEGs annotated as transcription factors involved in flavonoid metabolism

As many transcription factors (TFs) regulate the expression of flavonoid synthesis and transport, we also identified putative TFs based on the transcriptome. In our results, a total of 622 unigenes were determined as putative TFs (Table S4). The number of unigenes encoding MYB (128 DEGs) was the highest. Additionally, bHLH (77 DEGs), and WD40 (20) were identified. There is a protein complex that consists of MYB, bHLH and WD40, which plays a critical role in flavonoid metabolism. Moreover, ERF (79 DEGs), NAC (76 DEGs) and WRKY (69 DEGs) were found in this study, being the top abundant TFs. Most TFs were highly expressed in five samples, implying that they may take part in more metabolic processes of *R. roxburghii* compared with transporters (Fig. 3B),

### The P450 Family Gene expression

P450 enzymes involved in the generation of flavonoids [49]. By searching the DEGs, 221 DEGs were annotated as putative cytochrome P450 members and could be grouped into CYP subfamilies (Table S5). Some belong to the CYP 71 A, CYP 72 A and CYP 73 A. The others were annotated as CYP 81 A, CYP 86 A, CYP 90 A, CYP 98 B and so on. The CYP 98 family was reported to catalyze the 3-hydroxylation in the phenylpropanoid pathway [50]. Thus, the expression levels of cytochrome P450 would have a more significant effect on the biosynthesis of flavonoids in *R. roxburghii*. The expression pattern of the CYP genes was shown in this study (Fig. 3C). However, the deeper investigation of the roles of each CYP gene and the specific flavonoids still requires further study.

### Co-expression network analysis with WGCNA

To investigate the regulatory network and determine the key genes that influenced flavonoids in *R. roxburghii*. WGCNA was performed with the 1 310 non-redundant putative DEGs combining biosynthesis, transport, and regulators, leading to identifying of 19 distinct co-expression modules corresponding to clusters of correlated transcripts. The modules were labeled with different colors shown by the dendrogram (Fig. 4A), in which each tree branch constitutes a module. Each leaf of the branch was one unigene. The module eigengene can be considered a representative of the module’s gene expression profile [29]. Fig. 4B showed a hierarchical clustering dendrogram of the eigengenes. The total flavonoids, anthocyanin, and catechin contents acted as the trait data for a module-trait relationship analysis (Fig. 4C). The MEblack module (26 genes) presented the highest correlation with total flavonoids (r= 0.9, p= 5e-06), followed by the MEpurple (10 genes, r= 0.89, p=1e-05), MElightgreen (4 genes, r= 0.88, p= 1e-05) and MEblue (108 gene, r= 0.87, p= 3e-05). Besides, we noticed MEgreen (64 genes, r= 0.92, p= 1e-06) and MEyellow (78 genes, r= 0.83, p= 1e-04) displayed high correlation with anthocyanin and catechin accumulation, respectively. The genes in these modules were listed (Table S6). The results showed that different gene expressions played roles in specific flavonoids content.

**Fig.4.**
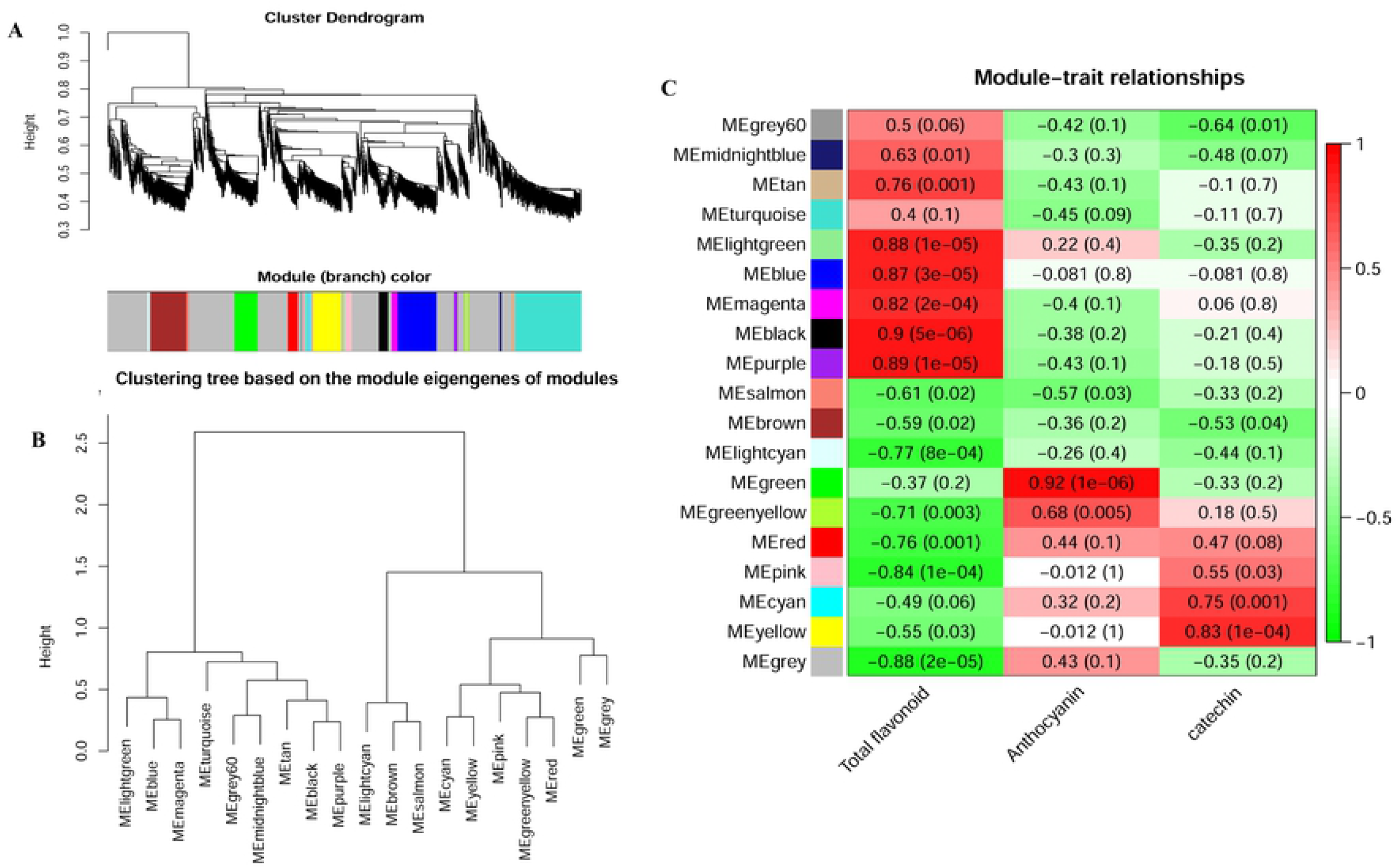
Weighted gene co-expression network analysis (WGCNA) of DEGs related to the flavonoid. (A) Gene dendrogram was obtained by clustering the dissimilarity based on consensus Topological Overlap with the corresponding module colors indicated by the color row. Each colored row represents a color-coded module that contains a group of highly connected genes. A total of 19 modules were identified. (B) Dendrogram of consensus module eigengenes obtained by WGCNA on the consensus correlation. The black line is the merging threshold, and groups of eigengenes below the threshold represent modules whose expression profiles should be merged due to their similarity. (C) Heatmap plot of consensus module eigengenes and flavonoid, anthocyanin and catechin content. The module type is shown on the left side. Numbers showed in the table report the correlations of the corresponding module eigengenes and tissue, with *p* values printed in the bracket. The table is color-coded by correlation according to the color legend. Intensity and direction of correlations are indicated on the right side of the heatmap (red, positively correlated; green, negatively correlated)

We selected the modules with correlation coefficient values ≥ 0.83 and analyzed eigengenes related to flavonoid biosynthesis (Fig. 5). The results showed that many biosynthesis enzymes play crucial roles in total flavonoids content (Fig.5A). Anthocyanin 3-O-glucosyltransferase and flavonoid 3’-monooxygenase (also named F3’H) were the two hub enzymes for anthocyanin content (Fig. 5B). Nevertheless, caffeic acid 3-O-methyltransferase, 4-coumarate-CoA ligase, and shikimate O-hydroxycinnamoyl transferase were essential for catechin accumulation (Fig. 5C). Interestingly, the number of P450 CYP749A22 and CYP72A219 was the highest (Fig.5A), implying that some subfamilies of cytochrome P450 were highlighted for flavonoids content.

**Fig. 5.**
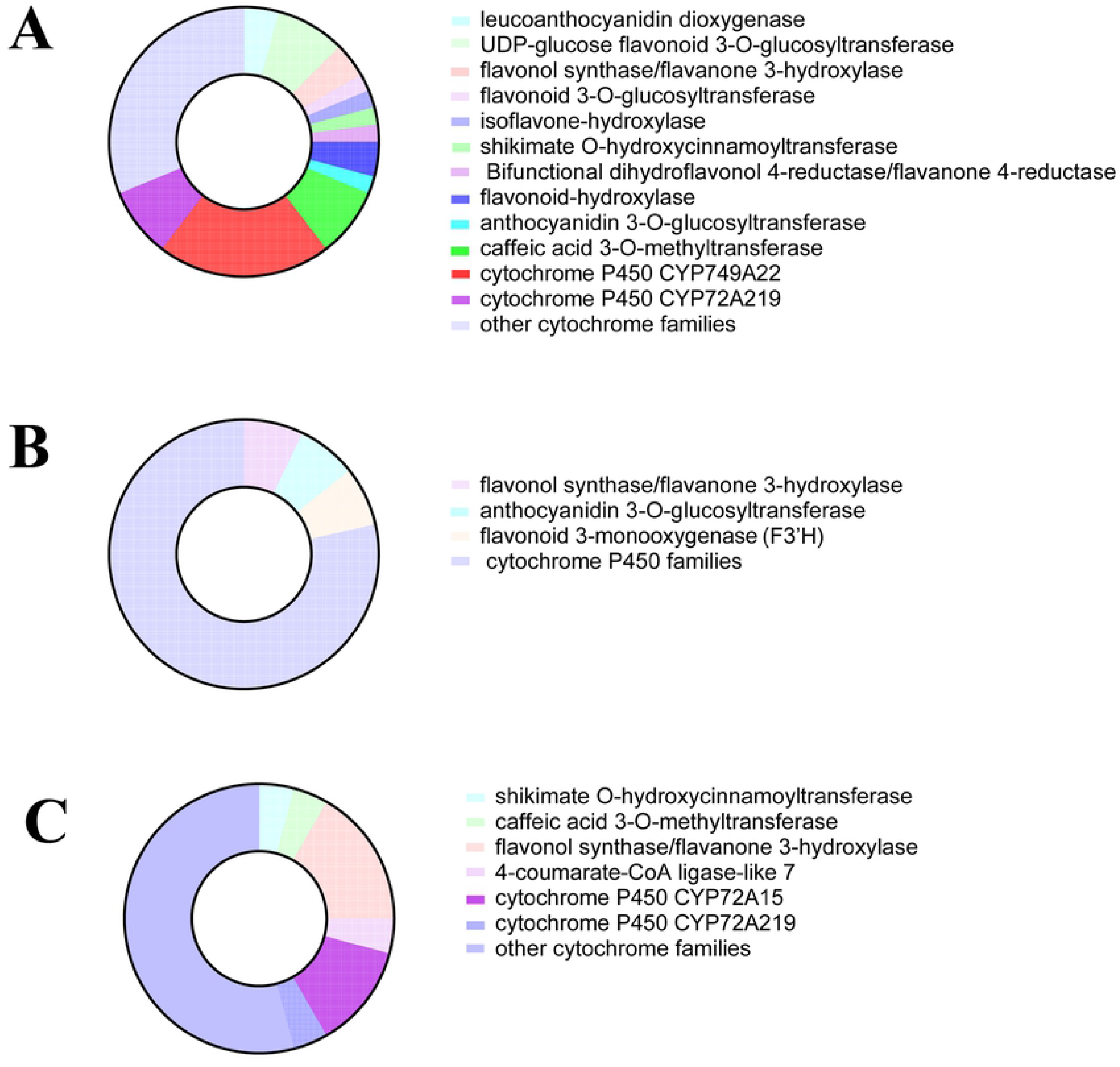
The genes related to flavonoid biosynthesis in module with correlation coefficient values ≥ 0.83. The genes had high connectivity with (A) flavonoids, (B) anthocyanin and (C) catechin content.

### Module visualization and hub genes

There were 26 eigengenes in the “black” module, including six flavonoid biosynthesis, 14 TFs and 6 transporters. To explore the critical genes that influences flavonoids content, we used Cytoscape software to visualize the network of the genes in “black” modules (Fig. 6A). Cytoscape representation of the 22 genes with WGCNA edge weight >0.80 indicated that these genes were highly positively connected in the “black” module. In the interaction network diagram, the outer layer consists of 12 genes related to TFs, including *RrWRKY45*, *RrTCP20* and *RrERF118*. In the middle of the network diagram, four flavonoid transporter genes were identified. The transcripts of 6 genes related to biosynthesis were identified in the network. All genes in the “black” module showed high expression in mature fruits based on transcriptome (Fig. 6B), further illustrating that the black module was very relevant to flavonoids content.

**Fig.6.**
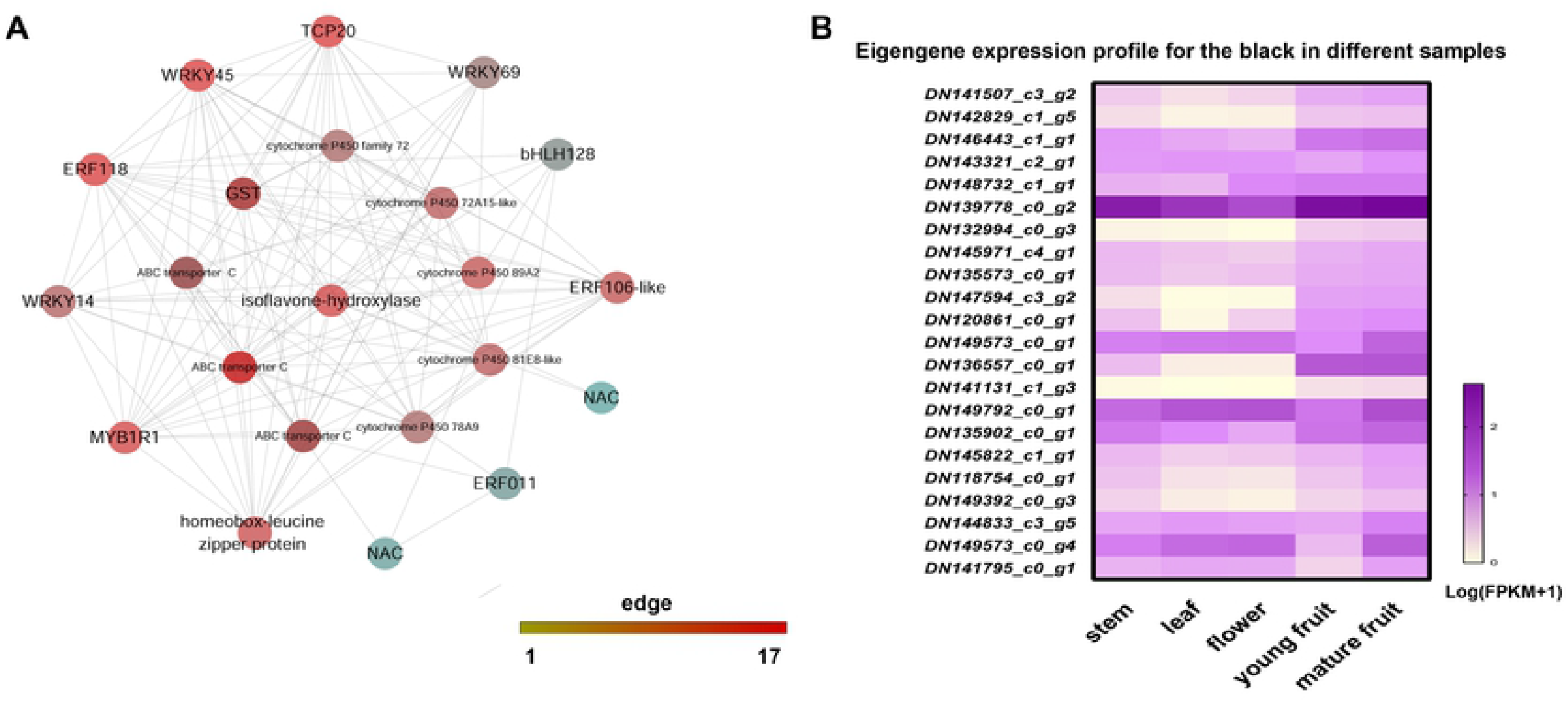
Cytoscape representation of co-expressed genes with edge weight ≥0.80 in module “black”. (A) The edge number of the genes ranges from 1 to 17 (colour-coded by the scale on the right from yellow through red). (B) Eigengene expression profile for the black in five samples.

### Confirmation of the hub genes using qRT-PCR

Based on ranking the connectivity of each node, we identified hub genes in the “black” module (edge ≥ 14). The expression levels of hub genes were determined by qRT-PCR using *β actin* as an internal control. We could obtain that the transcription levels of DEGs found in module ‘black’ were nearly consistent with the gene expression profiles obtained from RNA-seq (Fig. 7), indicating the reliability and accuracy of the RNA-seq analysis. Noticeably, the expression of most genes displayed lower in the stem, leaf, and flower while higher in fruits, especially mature fruits, confirming that the high correlation of hub genes with flavonoid accumulation. Additionally, the expression levels of the transcription factors *RrWRKY45* (*DN142829_c1_g5*), *RrTCP20* (*DN146443_c1_g1*) and *RrEFR118* (*DN141507_c2_g2*) were remarkably higher in fruits, implying that they exert functions on regulation flavonoids in *R. roxburghii*. Besides *cytochrome P450* (*DN136557_c0_g1*, *DN135573_c0_g1* and *DN145971_c4_g1*), *isoflavone-hydroxylase* (*DN143321_c3_g1*) was crucial for total flavonoids content.

**Fig. 7.**
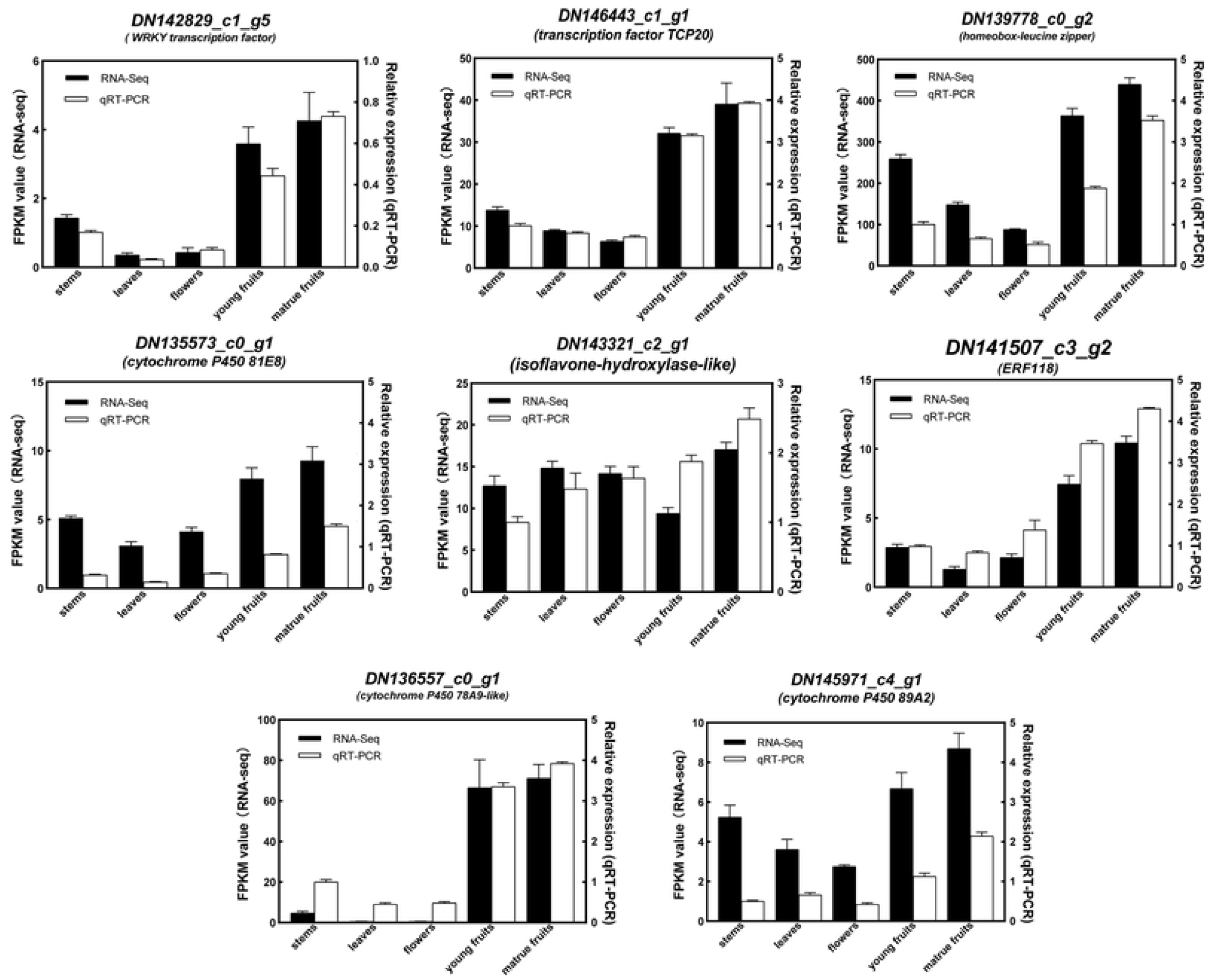
Comparison of expression profiles of nine hub genes from module ‘black’ as measured by RNA-seq and qRT-PCR. Columns represent expression determined by RNA-seq in FPKM values (left y-axis), while lines represent expression by qRT-PCR (right y-axis). The x-axis in each chart represents the five samples. Both for RNA-seq and qRT-PCR assay, the mean was calculated from three biological replicates. The Error bars show SD (standard deviation).

## Discussion

The transcriptome has become an effective tool to study biosynthesis mechanisms in plants. Transcriptome analysis for a flavonoid investigation in apple yellow fruit somatic mutation has been performed [34]. Another research reported candidate genes involved in flavonoid and stilbenoids metabolism *in Gnetum parvifolium* [36]. In the present study, we aimed to characterize the metabolic pathways of some important bioactive flavonoid compounds via a comprehensive in-depth investigation of the *R. roxburghii* transcriptome using RNA-seq. To generate data for an overview of the plant genetic composition, we performed samples for RNA preparation from different tissues of this species, and selected to decipher a comprehensive coverage of tissues as possible. Genomic survey sequencing for the genetic background of *R. roxburghii* was performed using next-generation sequencing technology by HiSeq 2500 sequencing in another study ^44^. The estimated genome size was 480.97 Mb based on the ratio of K-mer number to peak depth [51]. We obtained 63 727 unigenes in *R. roxburghii*, with an average GC content of 42.08% [35]. The coverage depth and accuracy of RNA-seq laid the foundation for obtaining gene expression. A total of 60 406 unigenes were annotated in at least one database [35]. Most annotations of these unigenes displayed highly similar strawberry (*Fragaria X ananassa*) [35].

*R. roxburghii* is rich in many flavonoid compounds [52]. Flavonoids extracted from *R. roxburghii* (FRR) leaves and fruits enhanced the radioprotective effect by inhibiting cell apoptosis [44]. However, flavonoids are complex and unevenly distributed in various tissues. Of these, anthocyanin is a typical kind of flavonoid compound, highly expressed in flowers [33]. The catechin content is relatively high among multiple flavonoid compounds in *R. roxburghii*, which was significant in leaves [5]. The flavonoids contents in mature fruits of *R. roxburghii* were higher than other tissues (Fig. 1). Therefore, they were selected as three trait data for module-trait relationship analysis.

We identified the 166 DEGs that encoded the known enzymes involved in the biosynthesis of the flavonoids and established a gene pool based on the *R. roxburghii* transcriptome. These DGEs showed different expression patterns in various tissues. For example, PAL catalyzes the first step in the biosynthesis of phenolic compounds [53]. High PAL activity was reported to be associated with the accumulation of anthocyanins in fruit tissues [54]. CHS is a critical enzyme in the flavonoid biosynthesis pathway [55]. Due to flavonoid composition mainly responsible for pollen tube growth and germination, and alterations in fruit color, the expression of CHS was increased [56, 57]. We analyzed the expression pattern of these DEGs related to *RrPAL* and *RrCHS* (Fig. 2). Strikingly, we found that most of them were plummet in reproductive organs (flowers and fruits) compared with the vegetative organs (leaves and stems). However, our results determined that the total content of flavonoids in fruits is significantly higher than in other organs. The reason may contribute to some critical genes that play more important roles in regulation of the flavonoid synthesis. Thus, it is essential to conduct a comprehensive and in-depth investigation of the *R. roxburghii* flavonoid metabolism.

Flavonoid accumulation may be a multifactorial process in the *R. roxburghii*, involving different strategies and the contribution of several transcriptional factors and transporters. Many transporters have been identified as critical factors in regulating flavonoid metabolisms, such as in *Lycium ruthenicum* Murray, *Dendrobium catenatum* and cranberry [14, 58, 59]. Therefore, we also analyzed candidate genes associated with transporters. Moreover, based on the analysis of putative DEGs involved in TFs, we revealed that the top four transcription factors were MYB (128), ERF (79), bHLH (77), and WRKY (68). Most of them were universally expressed genes among five samples. MYB, ERF and bHLH were all reported to regulate flavonoid biosynthesis in Arabidopsis, soybean, and grape [28, 60, 61]. These transcriptional factors may play essential roles in regulating genes associated with not only flavonoid biosynthesis, but also transporters, sequentially determining the flavonoid accumulation in *R. roxburghii* [14].

Overall, 1 310 DEGs were identified, considered to encompass the most relevant genes for flavonoids. WGCNA was implemented on transcriptomic data based on 1 310 DEGs. This approach constructed a network that segregated the expression profiles of 1 130 genes into 19 modules and showed high-quality gene clusters. We found different modules had connectivity with specific flavonoid compounds. We analyzed the eigengenes in the modules to evacuate the major factors which influenced the flavonoid accumulation (Fig. 5). Anthocyanin 3-O-glucosyltransferase and F3’H were the two hub enzymes for anthocyanin content. The previous reporters showed that the expression of anthocyanin 3-O-glucosyltransferase was significantly higher in black fruits than in white fruits of *Lycium ruthenicum* [33]. F3’H was hub genes in regulating anthocyanin of apple yellow fruit somatic mutation [34]. Overexpression of P450 promoted flavonoid biosynthesis [62]. The results showed that the expressions of cytochrome P450 CYP749A22 and 72A219, belong to heme-thiolate monooxygenase, were noticeably highlighted in regulating total flavonoids [63], indicating that they had close connectivity with flavonoids accumulation.

Hub genes displayed a high degree of connectivity in the module “black”, which showed a high correlation with flavonoids content based on the network. These putative genes were predicted to play more significant roles in total flavonoids accumulation in mature fruit. WRKY family not the only up-regulate the expression level of *CHS*, *F3*’*5*’*H*, and *F3*’*H* genes but also regulate the hydroxylation steps of flavonoids and MATE vacuolar transporter [64, 65]. *RrWRKY45* (*DN142829_c1_g5*) was confirmed to have high connectivity with flavonoids in the present study. In addition, TCPs regulate plant development and defense responses via stimulating flavonoid in Arabidopsis [66]. The potential anthocyanin regulator EFR118 was the hub gene in crabapple leaf color [67]. Our study presented that *RrTCP20* (*DN146443_c1_g1*) and *RrEFR118* (*DN141507_c2_g2*) were the hub genes in regulating flavonoid accumulation. The accumulation of total flavonoids and some specific compounds unevenly distributed in various tissues as it was stated in the results, which would eventually result in pharmaceutical differences. Thus, these findings could lead to genetic backgrounds involving biosynthesis, modification, transportation, and regulation of flavonoids. These findings provide a powerful genomic tool for the analysis of flavonoids regulatory network in the *R. roxburghii*.

The total flavonoids content was different and some specific compounds distributed in various tissues as it was stated in the results, which would eventually result in pharmaceutical differences. So, these findings could lead to genetic backgrounds involving biosynthesis, modification, transportation, and regulation of flavonoids. Flavonoid accumulation may be a multifactorial process in the *R. roxburghii*, involving different strategies and the contribution of several transcriptional factors and transporters. These findings provide a powerful genomic tool for the analysis of flavonoids regulatory network in the *R. roxburghii*.

## Acknowledgments

This work was supported by grants from the National Natural Science Foundation of China (Grant No. 31660554 and 31660046). The Joint Fund of the National Natural Science Foundation of China and the Karst Science Research Center of Guizhou province (Grant No. U1812401). Guizhou Educational project Qianjiaohe ([2021]309). The talent platform of Qiankehe ([2017] 5726-44). Sichuan Science and Technology Program (2021YJ0299 and 2021YFYZ0023). We would like to thank ISE (International science editing) for providing linguistic assistance during the preparation of this manuscript.

## Author contributions statement

X.H. and H.Y. designed the study. Z.W. and Y.J. analyzed the data. Y.Y. and Q.L. wrote the manuscript and conceived the experiments. All the authors approved the manuscript.

## Data availability

The RNA-Seq data is publicly available in NCBI GEO database under the accession number GSE122014.

## Supplementary Information

Table S1. Primers used for qRT-PCR analysis. (XLSX)

Table S2. The genes associated with flavonoid biosynthesis and their expression in five samples. (XLSX)

Table S3. The genes associated with transporters and their expression in five samples. (XLSX)

Table S4. The number of Candidate unigenes involved in transporters and transcription factors. (XLSX)

Table S5. The genes associated with transcription factors and their expression in five samples. (XLSX)

Table S6. The genes associated with cytochrome P450 and their expression in five samples. (XLSX)

Table S7. The list of all genes in “black”, “purple”, “lightgreen”, “blue”, “green” and “yellow” modules. (XLSX)

Fig. S1 Functional annotations of DEGs based on GO and KEGG enrichment analysis. (A) Different colors showed categories in the biological process (red), cellular component (green), and molecular functions (blue) relevant to plant physiology. One unigene may be matched to multiple GO terms. The left y-axis represents the numbers of unigenes with BLASTX matches to each GO term. The right y-axis indicated the percentage of genes. (B) Main functional categories were represented using various letters on the right. Metabolism (A); Environmental Information Processing (B); Genetic Information Processing (C); the Cellular Processes (D); and Organismal Systems (E). The X-axis represents the numbers of unigenes with BLASTX matches to each KEGG term. (TIF)

